# Forecasting histone methylation by Polycomb complexes with minute-scale precision

**DOI:** 10.1101/2023.06.16.545270

**Authors:** Moa J. Lundkvist, Ludvig Lizana, Yuri B. Schwartz

## Abstract

All animals use the Polycomb system to epigenetically repress developmental genes. The repression requires tri-methylation of Lysine 27 of histone H3 (H3K27me3) by Polycomb Repressive Complex 2 (PRC2), but the dynamics of this process is poorly understood. To bridge the gap, we developed a computational model that forecasts H3K27 methylation in *Drosophila* with high temporal resolution and spatial accuracy of contemporary experimental techniques. Using this model, we show that pools of methylated H3K27 in dividing cells are defined by the effective concentration of PRC2 and the replication frequency. We find that the allosteric stimulation by pre-existing H3K27me3 makes PRC2 better in methylating developmental genes as opposed to indiscriminate methylation throughout the genome. Applied to *Drosophila* development, our model argues that, in this organism, the intergenerationally inherited H3K27me3 does not “survive” rapid cycles of embryonic chromatin replication and is unlikely to transmit the memory of epigenetic repression to the offspring. We foresee that our model is adaptable to other organisms, including mice and humans.

## Introduction

All complex animals and plants use the Polycomb system to epigenetically repress developmental genes during cell differentiation (*1–4*). The repressed genes become enriched in histone H3 tri-methylated at Lysine 27 (H3K27me3) at their regulatory regions and open reading frames (*5–7*). H3K27me3 is required for the repression (*8*) and acts as a molecular mark that enables the repressed state after DNA replication (*9, 10*). The propagation of the repressed state is feasible because H3 molecules, methylated before the replication, are randomly partitioned between the two replicating chromatids (*11, 12*), which leads to the inheritance of the methylated chromatin state. Immediately after DNA replication, the density of the H3K27me3-containing nucleosomes is reduced by half to be fully restored before the next replication cycle.

In animal cells, the family of Polycomb Repressive Complexes 2 (PRC2) (*13, 14*) is responsible for mono-, di- and tri-methylation of H3K27. In flies and mammals, PRC2 methylates chromatin in two distinct modes (Figure 1). First, it randomly methylates H3K27 throughout the genome by the so-called hit-and-run mechanism (*13, 15–17*). Second, PRC2 methylates H3K27 within chromatin of specific loci where it is tethered by DNA elements, for example, mammalian CpG-islands (*15, 16*) or *Drosophila* Polycomb Response Elements (PREs) (*7, 18–21*). The latter correlates with a higher, nearly saturated, density of H3K27me3 nucleosomes around the tethering points (i.e. PREs) and is a hallmark of developmental genes repressed by Polycomb mechanisms. When methylating chromatin, PRC2 simultaneously engages with two histone H3 tails from different nucleosomes (*22*). The tail being methylated interacts with the catalytic centre of the complex while the other tail interacts with a non-catalytic subunit, called Esc (Extra sex combs) in *Drosophila* or EED (Embryonic ectoderm development) in mammals. If the Esc-interacting tail is already tri-methylated at K27, this allosterically stimulates the catalytic activity of PRC2 (*23*). Both hit-and-run and “tethered” methylation modes may benefit from the allosteric stimulation, but which of the two benefits most is currently unclear.

**Figure 1.**
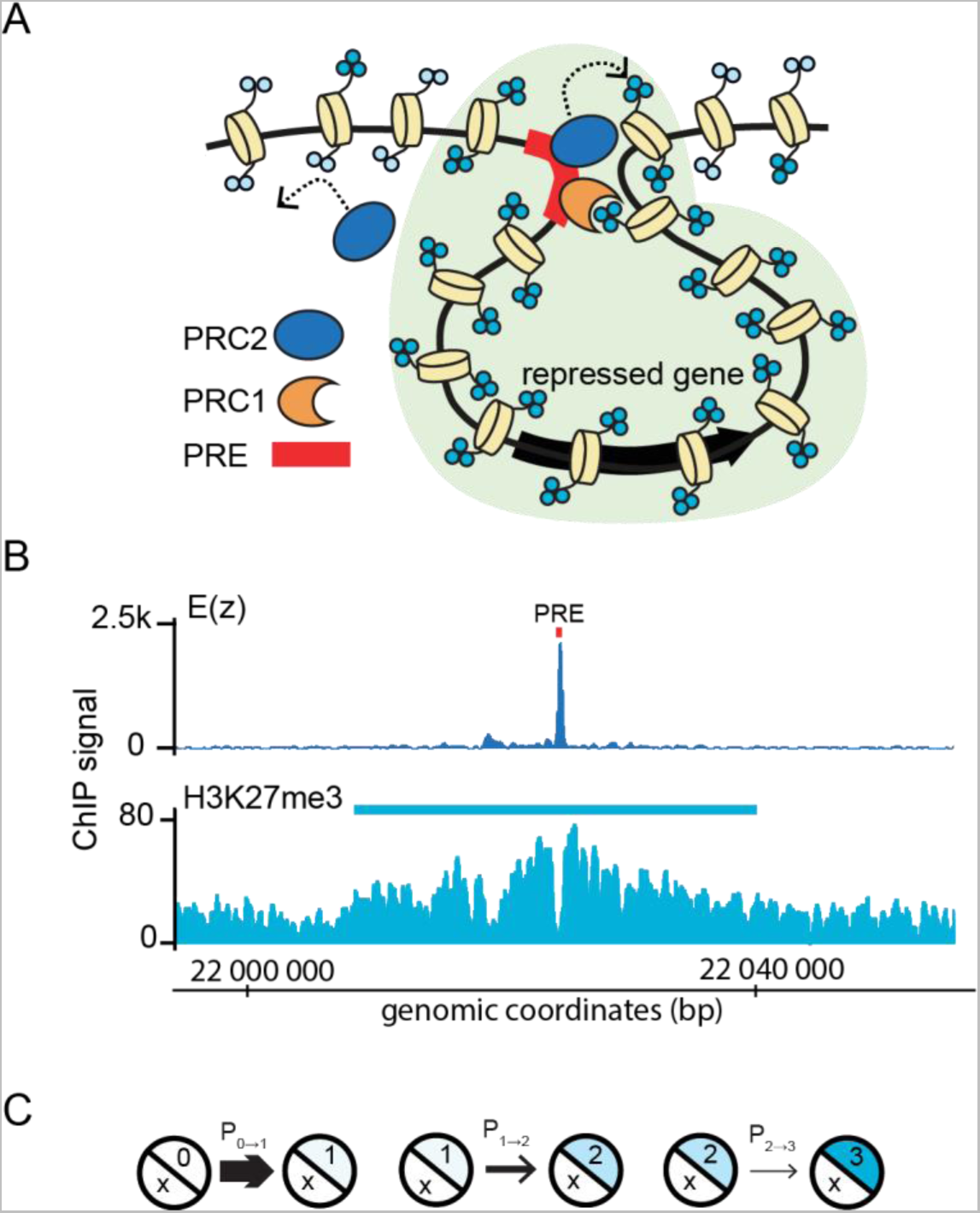
H3K27 methylation by PRC2. **A.** The schematic overview. PRC2 methylates H3K27 in two modes: by hit-and-run, which leads to bulk di- (light blue circles) and tri-methylation (dark blue circles), and by being bound to PREs. The latter leads to high levels of H3K27me3. PREs also bind Polycomb Repressive Complex 1 (PRC1), which interacts with H3K27me3 and stabilizes contacts between a PRE-bound PRC2 and methylated chromatin. **B.** An example of H3K27me3 profile around a PRE. The PRE (red box) is marked by strong ChIP-seq signal for the PRC2 subunit E(z). The blue box marks a region significantly enriched by H3K27me3 ChIP (corresponds to the shaded part in **A**). The scale is in dm3 genome release coordinates. **C.** Formal representation of the H3K27 methylation process. Each nucleosome has two H3K27 positions that can have 0-3 methyl groups attached. *In vitro* measurements indicate that the probability of adding a methyl group decreases for each methyl group already attached.

Although H3K27 methylation is essential for the epigenetic inheritance of the repression, little is known about the dynamics of the process, especially at specific developmental genes. For example, to what extent are the hit-and-run and tethered methylation modes coordinated? How long does it take to build up a sufficient density of tri-methylated H3K27 within genes repressed by the Polycomb system? What happens to the H3K27 “methyl mark” inherited from the parents? These questions, critical for understanding the epigenetic control of development, are still open.

Addressing these questions experimentally is challenging for several reasons. First, the methylation reaction is fast, non-processive, and has variable efficiency in adding the first, second, and third methyl group to K27 of the same H3 tail (*24, 25*). Second, the molecular mechanisms behind PRC2 tethering are not fully understood, which prevents the *in vitro* reconstitution of H3K27 methylation by the tethered complex. Finally, genetic approaches are limited because perturbations of the Polycomb system cause lethality during embryonic development.

To circumvent these issues, we developed a computational model that forecasts H3K27 methylation in *Drosophila* with high temporal precision. We show that a properly calibrated model of hit-and-run methylation can represent H3K27 methylation at PRE-equipped genes by converting the degree of PRC2 tethering into a locally increased methylation probability. Our model predicts that the allosteric stimulation of PRC2 by pre-existing H3K27me3 is necessary for broad and timely methylation of H3K27 around PREs. Applied to *Drosophila* embryonic development, the model forecasts that the intergenerationally inherited H3K27me3 does not “survive” rapid cycles of embryonic chromatin replication and, therefore, is unlikely to transmit the memory of epigenetic repression to the *Drosophila* offspring.

## Methods

### Algorithm and computational implementation

We represented the chromatin fibre as a one-dimensional vector of nucleosomes, where each nucleosome has two positions, corresponding to K27 of the two H3 molecules, which may have 0–3 methyl groups. Each nucleosome was set to contain 174 base pairs of DNA (*26*). Our computational algorithm changes the methylation state of each position stochastically according to specific methylation probabilities associated with e.g., PRC2 methylation reaction rates (also denoted catalytic rates). During each simulation round, the algorithm inspects all nucleosomes in random order and tries to add or remove one methyl group from every methylation position. Within a nucleosome the methylation positions are picked at random.

The model has 4 physical parameters that we use to calculate the methylation and demethylation probabilities: the base methylation probability (k_b_), the H3K27me3 stimulation (S_3_), the PRE tethering factor (F_PRE_), and the demethylation probability (k_D_). The other parameters we specify are the number of nucleosomes (N), the number of replicate simulations running in parallel (nGenomes), the total number of nucleosome inspection rounds (t_T_), and the number of nucleosome inspection rounds between replications (t_R_). t_R_ is adjustable but has a default setting for a 24-hour cell cycle. In this setting, the model performs t_R_ = 5760 inspection rounds per replication cycle, corresponding to one inspection round every 15 seconds. When calibrating the hit-and-run model to experimental data, we used nGenomes = 1000 and N=500. Calibration simulations ran for t_T_ = 57600 inspection rounds, which corresponds to 10 days in actual time. For simulations of specific gene loci, we used the same number of stretches (nGenomes = 1000) but varied N between 1000-2500, depending on the length of the genomic region. The full parameter settings for each figure are shown in Table S1.

To account for allosteric stimulation, we check the methylation status of a methylation position on a nucleosome other than the one being methylated. For simulations of the hit-and-run methylation, we choose one of the flanking nucleosomes. For locus-specific simulations, the partner nucleosome is chosen with a probability decaying as a power-law with the relative distance to the inspected nucleosome (for loop length l, p(l) ∼ l^-v^, where v = 0.75). After selecting the partner nucleosome, regardless of flanking or distant, we randomly choose one of the two methylation positions. If this position is tri-methylated, the allosteric stimulation is set to S_3_ (we calibrate S_3_ to experimental data). Otherwise, we set S_3_=1 (representing no stimulation).

Mathematically, we model the nucleosome-PRE contact probability using two critical observations. First, for distances shorter than a few thousand base pairs, the contact probability appears constant (*27*). We denote this distance L_PRE_. Second, over large distances, the contact probability decays as a power-law. This relationship appears in Hi-C experiments (*28*) and in theoretical polymer models (*29*). By bridging these two regimes, we define the effective contact function f(x), where x is the nucleosome-PRE distance in base pairs:

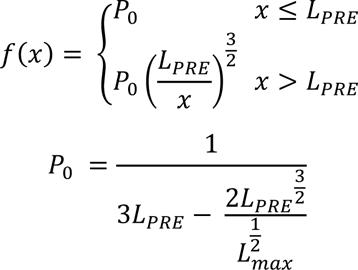

In the simulations, we use L_PRE_ = 2500 bp and L_max_ = 10^5^ bp, where L_max_ represents the distance beyond which the contact probability is zero. The decay exponent 3/2 indicates that we use a random (Gaussian) polymer model (*29*).

Experiments indicate that PREs have different abilities to tether PRC2. This notion comes from Chromatin Immunoprecipitation coupled to next-generation sequencing (ChIP-seq) showing that PREs are occupied by PRC2 to a different extent. We capture this in two model parameters: occupancy value (o_PRE,i_) and tethering factor (F_PRE_). While the tethering factor F_PRE_ controls the general stimulation effect of PREs, the occupancy value adjusts this effect individually for each PRE. The occupancy is calculated from ChIP-seq signals for the PRC2 subunit E(z) and normalized to vary between (0, 1]. Specifically, we calculated o_PRE,i_ as the sum of E(z) ChIP-seq reads from (*14*) within a 250bp window around the PRE, divided by the read sum for the PRE with the highest ChIP-seq signal. The model decides whether nucleosome methylation is stimulated by a PRE using a random number. If this number is less than the PRE contact probability, the nucleosome’s methylation probability increases with the PRE stimulation S_PRE_ = o_PRE,i_ * F_PRE_. If the nucleosome is not in contact with any PRE, there is no stimulation, and S_PRE_ = 1. When simulating chromatin methylation exclusively by hit-and-run action, the PRE stimulation routine is omitted.

Overall, the probability to acquire one methyl group is a product of several factors that must occur simultaneously. Apart from allosteric stimulation (S_3_) and the PRE stimulation (S_PRE_), we account for the decreasing probability of adding sequential methyl groups. The probability of adding the first methyl group to a position is 13k_b_, the probability of adding the second group is 4k_b_, and the probability of adding the third group is k_b_ (*25*). In summary, we define the methylation probability of a position as: k * S_3_ * S_PRE_, where k is either 13k_b_ or 4k_b_ or k_b_.

Nucleosomes can also lose methyl groups. In our implementation, this happens in two ways. First, we remove one methyl group with a probability k_D_, which represents enzymatic demethylation or histone exchange. Second, all nucleosomes have a 50% chance to become unmethylated during replication, which happens every t_R_ inspection round.

All simulations were implemented as JavaScript code and run with the node.js environment using the built-in modules fs and zlib, and the Papa Parse package. Simulation outputs were written to json files. The plotting and analysis were performed in Python. The model implementation is freely available at https://github.com/lizanalab/forecasting2023lundkvist.

### Simulation outputs and data comparison

#### Definition of forecasted H3K27 methylation profiles and H3K27me3-enriched regions

The H3K27 methylation profile for a simulated string of nucleosomes was defined for each of the four possible methylation states (un-, mono-, di-, and tri-methylated). The profile is the frequency of each nucleosome to be in that state over a set of runs with the same settings. As each nucleosome has two H3K27, the possible value per nucleosome for each state varies between 0 and 2. The profile was defined as the mean from five selected inspection rounds over a cell cycle for 1000 simulated stretches. The inspection rounds 0, 1440, 2880, 4320, and 5759, within a cell cycle with a total of 5760 inspection rounds, were selected. These inspection rounds correspond to time points immediately after replication, immediately before the next replication, and three evenly spaced points in between.

The H3K27me3-enriched regions were defined as stretches of nucleosomes within PRE-equipped loci where the profile was 3 Standard Deviations (SD) above the mean profile from simulations that did not include PREs (hit-and-run methylation only), for at least three nucleosomes in a row. The H3K27me3-enriched regions separated by less than 5kb were merged.

To define the PRE-equipped loci, computationally defined PREs from (*14*) were divided into groups where PREs within 100kb of each other marked the same locus. Only the PRE-equipped loci enriched for H3K27me3 in ChIP-seq experiments performed with chromatin from Kc167 or EZ2-2 cells (total of 97) were used for comparison with experimental measurements (see below).

There are some cell line-specific differences among the 97 loci. First, not all loci are repressed in both cell lines. Of the 97 modelled loci, 89 were defined in the Kc167 data set, and 92 in EZ2-2 (one Kc167 locus became two in EZ2-2, since the long tri-methylated region was interrupted in EZ2-2). The specific parameters used for simulating the set of loci in each of the two cell lines are shown in Tables S2 and S3. Furthermore, among the loci that are repressed in both cell lines, the transcriptional activity of surrounding genes sometimes differs, which affects the extent of tri-methylated regions. Seven loci were excluded from the experiment-to-experiment comparison in Figure 3D for that reason.

#### Definition of transcriptionally active regions

The RNA-seq data from (*14, 30*) SRR8743273, SRR1573098 were aligned to the Drosophila genome in dm3 release coordinates using bowtie2 (Langmead and Salzberg, 2012) with the following parameters: `bowtie2 -- phred33 -p 8 -x ../Dm3_bowtie_index_files/Dm3 -1 reads1.fastq.gz -2 reads2.fastq.gz.` The aligned reads were filtered and sorted with samtools: `samtools view -q 30 -b aligned_reads.bam` and samtools sort -@ 8 aligned_reads_q30.bam, and the profile generated with bedtools`: bedtools genomecov -ibam aligned_reads_q30_sorted.bam -bg > profile.bdg.` The profiles were scaled by their millions of mapped reads.

A genomic region was defined as transcriptionally active if the score was above the 0.25th quantile. The genomic coordinates of the regions were converted to the closest nucleosome indices within corresponding simulated loci. Since active genes are not included in the model, transcriptionally active regions are used to cut off the simulated me3 regions. If a me3 region that is interrupted by an active region has PREs on the other side of the active region, the me3 region continues on that side. If not, that part of the me3 region is lost as well.

#### H3K27me3 ChIP-seq data processing and definition of enriched regions.

Two H3K27me3 ChIP-seq experiments, with two replicates each, were used: one performed with chromatin from Kc167 cells (Cubenãs-Potts et al., 2017) (GEO IDs: SRR3452731, SRR3452732), and another one performed with chromatin from EZ2-2 cells grown at 25°C (permissive temperature) (Lee et al., 2015) (GEO IDs: SRX699107, SRX811237). The corresponding sequencing reads were aligned to the *Drosophila* genome in dm3 release coordinates using bowtie2 (*31*) with the following parameters: `bowtie2 --phred33 -p 8 -x ../Dm3_bowtie_index_files/Dm3 -1 reads1.fastq.gz -2 reads2.fastq.gz`. The aligned reads were filtered with samtools: samtools view -h -b -@ 8 -q 30 -o filtered_aligned_reads.bam and H3K27me3 profiles generated using MACS2 (*32*): `macs2 pileup -i filtered_aligned_reads.bam -o output.bdg -f BAM -- extsize 180`

Regions with H3K27me3 ChIP-seq signal 3 SD above that within generic transcriptionally inactive chromatin (defined as BLACK in Kc167 cells by (*33*)) were considered enriched. A H3K27me3-enriched region was set to begin when ChIP-seq signal remained above the 3 SD threshold for at least 15bp, and it was set to end when the signal remained below 3 SD for as long. The H3K27me3 regions closer than 5kb were merged. To compare with regions forecasted by the model, the start and stop coordinates of the experimentally defined H3K27me3-enriched regions were converted to nucleosome indices of the corresponding locus based on the distance to the closest PRE.

## Results

To understand the quantitative properties of the epigenetic memory based on H3K27 methylation, we built a computer model that forecasts histone methylation by PRC2 at *Drosophila* genes regulated by PREs. We wished to have a model with minute-scale precision and spatial accuracy that matches contemporary experimental methods such as Chromatin Immunoprecipitation coupled to massively parallel sequencing (ChIP-seq).

We envisioned a model where chromatin was represented as a vector of nucleosomes each having two H3K27 methylation positions. The model would periodically inspect all positions and record whether they have zero, one, two, or three methyl groups. Between inspections, each H3K27 could acquire or lose one methyl group or experience no change. Such a model can be implemented as a computational Monte-Carlo simulation, provided that realistic probabilities of each outcome (denoted transition probabilities) are derived from experimental data.

Unfortunately, the methylation reaction by the PRE-tethered PRC2 has not yet been reconstructed *in vitro*, and its biochemical characteristics are largely unknown. Therefore, realistic transition probabilities for this reaction are difficult to estimate. In contrast, methylation by the reconstituted PRC2 in solution was extensively studied *in vitro* (*24, 25*). This reaction is mechanistically similar to the hit-and-run methylation of H3K27 by diffusing PRC2. As we discuss in detail in the following section, in dividing cells, the bulk of mono- di- and tri-methylated H3K27 is produced by hit-and-run methylation. We hypothesized that it might be possible to first construct a realistic computational Monte-Carlo model of the hit-and-run methylation and then adapt it to represent H3K27 methylation by tethered PRC2. We could achieve this by converting the degree of PRC2 tethering by PREs into an increased methylation probability.

### Monte-Carlo simulation of the hit-and-run methylation by PRC2

To simulate the hit-and-run methylation, we calculated the associated transition probabilities using a set of computations and logical decisions (Figure 2A). We cycled through this set of decisions/computations for each H3K27 methylation position within the simulated chromatin fibre during every inspection round. Below, we briefly describe some of these decisions.

**Figure 2.**
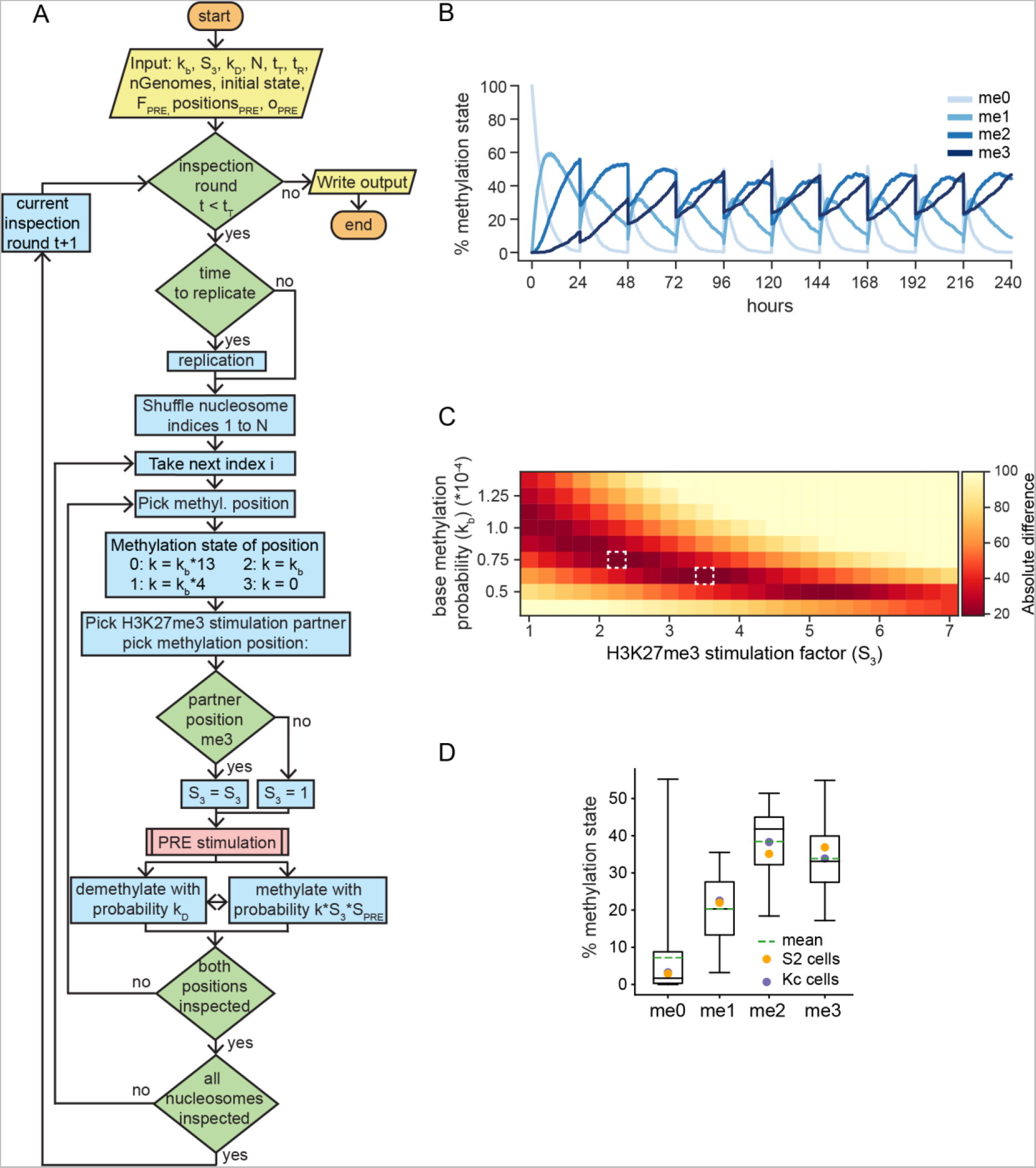
Hit-and-run methylation model. **A.** The schematic of the simulation algorithm. Orange ovals indicate the algorithm start and stop, yellow parallelograms show the input and output, blue boxes correspond to processes, the pink box indicate the “tethered-methylation” subprocess and green diamonds represent logical decisions (see Methods for further explanations and description of input parameters). **B.** The dynamics of methylation pools from a representative hit-and-run simulation under optimized parameters. Here and in **C** simulations started with completely unmethylated nucleosomes. **C.** Optimization of base methylation probability (k_b_) and H3K27me3 stimulation (S_3_) parameters. The heat-map representation of the sums of absolute differences between the mean fractions of un-, mono-, di-, and tri-methylated H3K27 forecasted by the model and those measured by mass-spectrometry in S2 and Kc cells indicates that parameter settings S_3_ = 3.5; k_b_ = 0.625×10^-4^ and S_3_ = 2.25 and k_b_ = 0.75×10^-4^ (marked with dashed white boxes) result in predictions most closely resembling the experimental measurements. The fractions of variously methylated H3K27 positions predicted by the model were calculated based on 10 inspection rounds from 1000 different simulations. The inspection rounds were selected randomly between the 216^th^ and 240^th^ simulation hour (10^th^ cell cycle) when the system has reached the steady state. **D.** Side-by-side comparison of forecasted and measured fractions of variously methylated H3K27 molecules. Box plots show the percentage of methylation positions in each state for S_3_ = 3.5 and k_b_ = 0.625×10^-4^ calculated as above. Here and in all subsequent figures the box plots indicate the medians (solid lines), averages (dashed lines) and span interquartile range with whiskers covering the full extent of the data. The average percentages of corresponding methylated H3K27 molecules measured by mass-spectrometry are shown with yellow (S2 cells) and purple (Kc167 cells) dots.

First, from *in vitro* reaction with soluble PRC2, we know that the complex is progressively less efficient in adding the second and the third methyl groups to already methylated H3K27. Therefore, the model evaluates the methylation state of each methylation position and adjusts the corresponding transition probability such that the ratios between transition probabilities P_0→1_: P_1→2_: P_2→3_ match the efficiencies of the corresponding reactions measured as 13:4:1 (*25*) (Figure 1C). These numbers relate to typical enzymatic reaction parameters 𝑘_𝑐𝑎𝑡_ and 𝐾_𝑀_, where 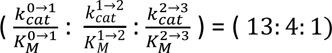

Second, the catalytic activity of PRC2 is stimulated by its own product, H3K27me3 (*23, 34*). Mechanistically, this is achieved when PRC2 bridges two nucleosomes and “senses” the methylation state of one of them while methylating the other (*35*). We denote this feature H3K27me3 stimulation and model it by checking the state of one of the methylation positions on a flanking nucleosome (picked at random). If the position is tri-methylated, all transition probabilities increase by a factor S_3_.

The model must inspect nucleosomes reasonably often. If the time between inspections is too long, some H3K27 could acquire more than one methyl group. We derived the required inspection frequency from catalytic properties of PRC2, which adds one methyl group per catalytic cycle (*25*). When adding the first methyl group to unmethylated H3K27, the PRC2 turnover rate *in vitro* was estimated as 0.014± 0.001 min^-1^ (*25*), 0.017± 0.001 min^-1^ (*36*) or 1.01 ± 0.39 min^-1^ (*24*). From these values, we concluded that if H3K27 positions are inspected every minute or more frequently, at most one methyl group would be added.

Monte-Carlo simulations do not have a built-in time scale. To circumvent this problem, we related the inspection frequency to the cell cycle length. In culture, typical *Drosophila* cells replicate their genomes and divide approximately every 24 hours (i.e., every 1440 minutes) (*37*). Therefore, we added periodic DNA replication events to the model and set the number of inspection rounds between replications to 5760 (i.e., every 15 seconds). When DNA replicates, parental nucleosomes randomly redistribute between the two daughter genomes, and newly synthesized unmodified histones fill the gaps. To recapitulate this process, with 50% probability, the model converts the H3K27 positions on nucleosomes to the unmethylated state every 5760^th^ inspection round.

Besides replication, methyl groups can be removed by specific histone demethylases (*38–40*). While the model accounts for this possibility (Figure 2A), experiments with PRC2 knock-out cells (*17*) indicate that untargeted demethylation is much slower compared to the dilution following DNA replication. Therefore, we set the demethylation probability to 0 when simulating hit-and-run methylation in dividing cells. We evaluate the role of H3K27 demethylation in the non-dividing cells in a later section.

The simple model described above forecasts methylation by untethered PRC2 in cultured *Drosophila* cells. Two lines of evidence argue that most of the mono-, di- and tri-methylated H3K27 in dividing cultured cells is produced by untethered PRC2. First, genomic mapping indicates that H3 molecules mono-, di- and tri-methylated at K27 are scattered throughout the genome and not limited to regions around PREs where PRC2 is tethered (*13, 17*). Second, Feller and co-authors used quantitative mass-spectrometry to estimate that, in cultured Drosophila cells, 33.9-36.9% of all histone H3 molecules are tri-methylated at K27. (*41*). The regions around *Drosophila* PREs that are highly enriched in H3K27me3 (*7, 21, 42*) have a cumulative length of 9.4Mb (*21*), which corresponds to 5.2% of the 180Mb-long genome (*43*). Even if all H3 molecules within these highly enriched regions are tri- methylated at K27, this accounts only for one-seventh of all H3K27me3, indicating that the bulk of the tri-methylated H3K27 is scattered throughout the inactive genome and produced by hit-and-run action of PRC2.

If most of the methylated H3K27 in dividing cultured cells is produced by untethered PRC2, we can calibrate critical parameters of our model using mass-spectrometry measurements of bulk levels of methylated H3K27 in cultured *Drosophila* cells (*41*). To this end, we ran a series of simulations varying the base methylation probability k_b_ and different extents of H3K27me3 stimulation (S_3_). We found that the relative levels of H3K27me0, H3K27me1, H3K27me2, and H3K27me3 fluctuate throughout the cell cycle over a wide range of k_b_, where H3K27me3 grows slowly until being diluted two-fold by DNA replication (Figure 2B). We sampled the simulated data at random points over the cell cycle to mimic the mass-spectrometry measurements in an un-synchronized cell culture and calculated the average fraction of variously methylated H3K27 using 10 random inspection rounds per simulation over 1000 runs. Figure 2C illustrates how the simulation results deviate from experimental data at a range of parameter values for S_3_ and k_b_. In 500 independent calculations, two parameter pairs gave the best match with experimental data: S_3_ = 3.5; k_b_ = 0.625×10^-4^ and S_3_ = 2.25; k_b_ = 0.75×10^-4^. The former gave the best fit 33.4% of the time, and the latter 49.8% of the time. As S3 = 2.25 falls outside measured H3K27me3 stimulation values, we chose S_3_ = 3.5; k_b_ = 0.625×10^-4^ parameter setting for further simulations. As illustrated by Figure 2D, this setting recapitulates experimental measurements, and we conclude that our Monte-Carlo model accurately forecasts hit-and-run methylation by PRC2.

### Accounting for tethered PRC2 recapitulates H3K27 methylation at PRE-equipped genes

With the hit-and-run methylation model at hand, we wished to explore if such a model could be adapted to recapitulate the H3K27 methylation at PRE-equipped genes by converting the degree of PRC2 tethering by PREs into a higher methylation probability. Thus, we supplemented our algorithm with a new module (Figures 3A, 2A) that decides if the nucleosome is engaged with a PRE-tethered PRC2. If not, the transition probability remains identical to the hit-and-run methylation model. We assume that the PRE-tethered PRC2 retains the same P_0→1_: P_1→2_: P_2→3_ ratios as the untethered PRC2, but that the tethering increases the chance of methylating the nucleosome. We describe this increase with a parameter denoted tethering factor (F_PRE_). As PREs differ in their ability to retain PRC2, we further adjust the degree of tethering by individual PREs with a PRE-specific occupancy parameter (o_PRE,I_, see Methods for details).

The above defines the methylation capabilities of the PRE-tethered PRC2. But what is the likelihood for a nucleosome to be contacted by a PRE-tethered PRC2 complex? Theoretical polymer models, like the Gaussian chain or the fractal globule model (*29*), and chromatin conformation (Hi-C) measurements (*28*) suggest that, over long distances, the contact probability between any two genomic regions decays with distance as a power-law. However, over short distances (below 5kb), such estimate is unreliable as the Hi-C resolution is limited by the size of DNA fragments generated after digesting the chromatin with a restriction endonuclease. Moreover, the chromatin of genes repressed by the Polycomb system displays an unusually high degree of intermixing, suggesting that the probability of contacts in the immediate vicinity of PREs is largely distance-independent (*27*). To reconcile these two regimes, we modelled the contact probability as constant over a short region (2.5kb on each side of a PRE) and decaying as a power law for longer distances.

Like in the hit-and-run model, PRE-tethered methylation also considers H3K27me3 stimulation (embedded in the parameter S_3_). When modelling hit-and-run methylation, we considered nearest neighbour interactions. However, two nucleosomes may be far apart along the DNA and brought in proximity via chromatin looping (*35*). Therefore, for PRE-equipped genes, we picked partner nucleosome distances from a power-law distribution.

To evaluate the fidelity of our amended H3K27 methylation model, we simulated 97 *Drosophila* loci equipped with one to fourteen PREs and compared the simulated profiles of tri-methylated H3K27 to those derived from ChIP-seq data. To this end, we used four profiles from two ChIP-seq experiments with unperturbed Kc167 and EZ2-2 cultured cell lines, each with two replicates (*17, 44*). A well-performing model should forecast the methylation consistently, such that the outcomes of two independent simulations differ no more than two replicate ChIP-seq experiments. Moreover, the model should produce H3K27me3 distributions that match those from ChIP-seq assays at least as closely as two independent experimental distributions. As illustrated by the comparison of predicted and experimentally defined H3K27me3-enriched regions within the *bithorax* complex (the cluster of homeotic genes *Ubx*, *abd-A* and *Abd-B*) (Figure 3B), this is indeed the case at some loci. However, at others, for example the *earmuff* (*erm*) locus, the predicted and experimentally defined H3K27me3 distributions differ substantially (Figure 3C). Here, the experimental H3K27me3 profile drops to zero immediately to the right of the PREs in contrast to the simulated distribution. This drop coincides with a cluster of transcriptionally active genes (*17, 30*) (Figures 3C). Transcriptional activity is known to inhibit PRC2-mediated H3K27 methylation by means that that are not fully understood (*7, 45, 46*) and, therefore, difficult to model explicitly. To enable fair comparison and account for inhibitory effect of transcription, we defined transcriptionally active regions from RNA-seq data (see Materials and methods for details) and introduced a rule that “stops” an H3K27me3-enriched region when bordered by a transcriptionally active gene.

**Figure 3.**
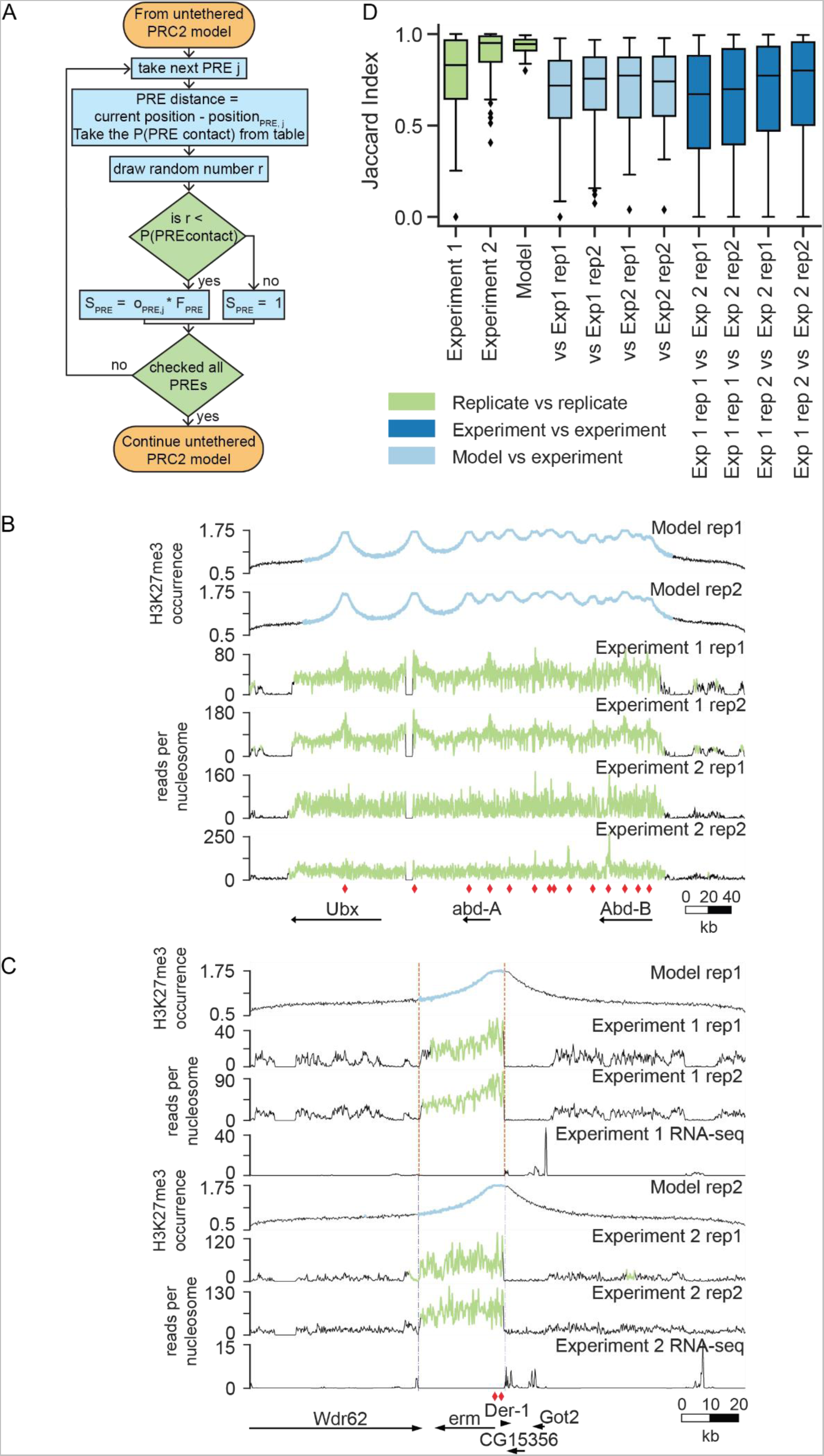
Modelling H3K27 methylation at PRE-equipped genes. **A.** The schematic of “tethered methylation” subprocess of the simulation algorithm. During each inspection it checks whether a nucleosome is engaged with a PRE-tethered PRC2 and increases the methylation probability accordingly. **B.** The H3K27me3 distribution over the bithorax complex forecasted by the model is juxtaposed with distributions derived from two different ChIP-seq experiments. Here and in **C**, the simulated profiles show mean values from 5 inspections within one cell cycle over 1000 parallel simulations. The profiles derived from experimental measurements show mean ChIP-seq reads over a nucleosome **C.** The H3K27me3 and RNA-seq profiles over the *erm* locus. The dotted lines mark the edges of transcriptionally active regions. **D.** Box plots of Jaccard indices comparing H3K27me3-enriched regions defined from experimental and simulation replicates (green), individual simulations and experiments (light blue) and between different experiments (dark blue). Note that simulations are less noisy and identify H3K27me3-enriched regions as well as ChIP-seq assays.

To quantify the comparison between H3K27me3-enriched regions we used the Jaccard index (*J*). This is a statistic used to gauge the similarity between two objects consisting of multiple binary attributes (*47*). Customary in the network science and image analysis, the Jaccard index is calculated by dividing the intersection between the two objects by their union. This way, two identical regions enriched in H3K27me3 have *J*=1 while two completely non-overlapping regions have *J*=0. Figure 3D shows boxplots of the Jaccard indices calculated from pairwise comparisons of H3K27 methylated regions across the 97 PRE-equipped *Drosophila* loci, defined empirically or by simulation, and accounting for transcriptional activity. Overall, it indicates that our simulations forecast the H3K27 tri-methylated regions with a consistency that matches that of the two independent ChIP-seq experiments. This result suggests that it is possible to accurately recapitulate H3K27 methylation at PRE-equipped genes by converting the degree of PRC2 tethering by PREs into an increase in the methylation probability.

### The role of PRC2 stimulation by H3K27me3

The stimulation by pre-existing H3K27me3 is an evolutionarily conserved feature of PRC2. When first discovered, it was hypothesized to maintain H3K27 methylation at genes repressed by the Polycomb system and restore the methylation in daughter cells (*23*). Taken to the extreme, this view suggests that PRC2 tethering is only needed to establish the high degree of H3K27 tri-methylation within a region, while the allosteric stimulation of untethered PRC2 suffices to renew it after DNA replication. This extreme view is at odds with transgenic experiments, which indicate that tri-methylated H3K27 in dividing cells is gradually lost from a transgene after the excision of the PRE DNA (*9, 10*). Simulations confirm that propagation of elevated H3K27me3 in replicating chromatin requires PREs to tether PRC2. As illustrated by modelling the *H15* locus, the level of H3K27me3 coincides with the hit-and-run prediction after about five replication cycles when all three PREs of the locus are removed (Figure 4A).

**Figure 4.**
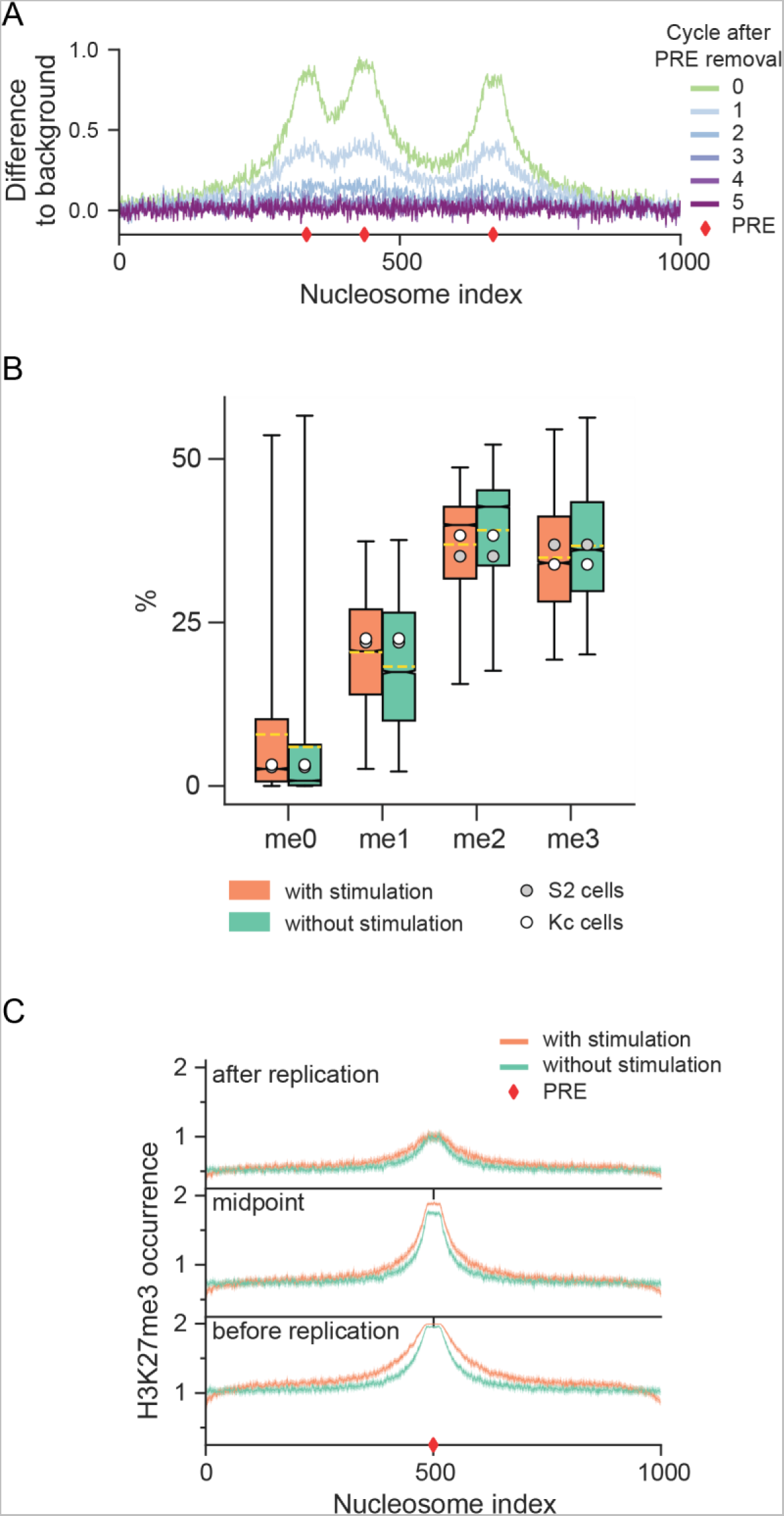
Contributions of PRC2 stimulation by H3K27me3. **A.** The simulated H3K37me3 profiles over the *H15* locus during five cell cycles after removing PREs. Shown are the differences in the H3K27me3 occurrence relative to a chromatin fibre without PREs. The occurrence was calculated from 5 inspections within one cell cycle over 1000 parallel simulations. **B**. Box plots showing the percentage of H3K27 positions in each methylation state from 10 random inspections of 1000 different simulations (when they have reached the steady state) performed with (orange, S_3_ = 3.5 and k_b_ = 0.625×10^-4^) or without (green, S_3_ = 1 and k_b_ = 1.125×10^-4^) H3K27me3 stimulation. The means are shown as yellow dotted lines. The average percentages of corresponding methylated H3K27 molecules measured by mass-spectrometry are shown for comparison. **C.** The mean H3K37me3 profiles over the *trachealess* (*trh*) locus simulated with (orange line) and without (green line) H3K27me3 stimulation captured during the first, the middle, and the last inspections of the cell cycle. The 95% confidence interval from 1000 simulations are shown as shaded areas.

If the stimulation by pre-existing H3K27me3 is not enough to propagate the methyl mark, what is it good for? When optimizing parameters for hit-and-run methylation, the best match to experimental bulk levels of mono- di- and tri-methylated H3K27 is achieved with some degree of allosteric stimulation (S_3_ = 2.25 or S_3_ = 3.5, Figure 2C). However, a less optimal but good fit is also achievable without it (S_3_ = 1, Figure 2C). This requires a higher level of the base methylation rate (k_b_ = 1.125×10^-4^). Conceivably, cells can attain a high methylation rate by increasing the PRC2 production or by incorporating the auxiliary subunit Jarid2 into the complex. Jarid2 contains a H3K27 mimic that can be methylated by PRC2’s catalytic core and stimulate PRC2 activity (*48*).

However, compensating by a higher base methylation probability is not perfect (Figure 4B, 1S). While the methylation values are similar, this new stimulation-free parameter setting skews the ratio between H3K27me1: H3K27me2: H3K27me3 towards higher methylation states compared to that measured experimentally or produced by simulations that account for the allosteric stimulation (S_3_ = 3.5; k_b_ = 0.625×10^-4^). Still, the differences are small and therefore unlikely to explain the evolutionary pressure to have PRC2 stimulation by H3K27me3.

An alternative hypothesis suggests that allosteric stimulation is required for the H3K27me3 spreading around tethered PRC2. Supporting this hypothesis, the H3K27-enriched regions around tethered PRC2 became shorter in mouse embryonic stem cells with mutated PRC2 insensitive to the allosteric stimulation (*49*). Unfortunately, the interpretation of that result is confounded by notably weaker binding of the mutated PRC2 (*49*). To investigate the spreading hypothesis, we simulated H3K27 methylation around the single PRE of the *trachealess* (*trh*) locus under the best fitting parameters that either include or exclude the allosteric stimulation. We focused on the H3K27me3 distributions immediately after DNA replication, in the middle of the cell cycle, and just before DNA replication. As illustrated by Figure 4C, in the presence of allosteric stimulation, the regions enriched in H3K27me3 are broader at all three time points. Furthermore, the H3K27me3 is regained faster and reaches saturation in the vicinity of the PRE well before DNA replication.

To summarize, our simulations support the model where the allosteric stimulation of PRC2 by H3K27me3 enables a broad H3K27 tri-methylation profile around PREs. They also suggest that it hastens the recovery of H3K27me3 there, so that the required level of tri-methylation is reached before the next replication event.

### Differentiated cells require H3K27 demethylation

In mitotically dividing cells, the pool of methylated H3K27 is regularly diluted by the influx of unmodified H3 molecules due to chromatin replication. Methyl groups can also be removed from H3K27 by histone demethylating enzymes (*38, 40, 50, 51*), and nucleosomes are lost and replaced during transcription (*52*). We will collectively refer to these replication-independent processes as H3K27 demethylation.

As shown above, in dividing cells, the H3K27 methylation is accurately forecasted without explicit accounting for H3K27 demethylation. This is true for both hit-and-run methylation of the transcriptionally inactive chromatin as well as tethered methylation within PRE-equipped loci. In fact, the simultaneous optimization of the base methylation probability (k_b_) and the demethylation probability (k_D_) parameters shows that k_D_ = 0 produces the best hit-and-run methylation forecast (Figure 5A). This is consistent with observations in cultured *Drosophila* cells, where the replication-independent loss of H3K27 methylation across the genome is slow (*17*).

**Figure 5.**
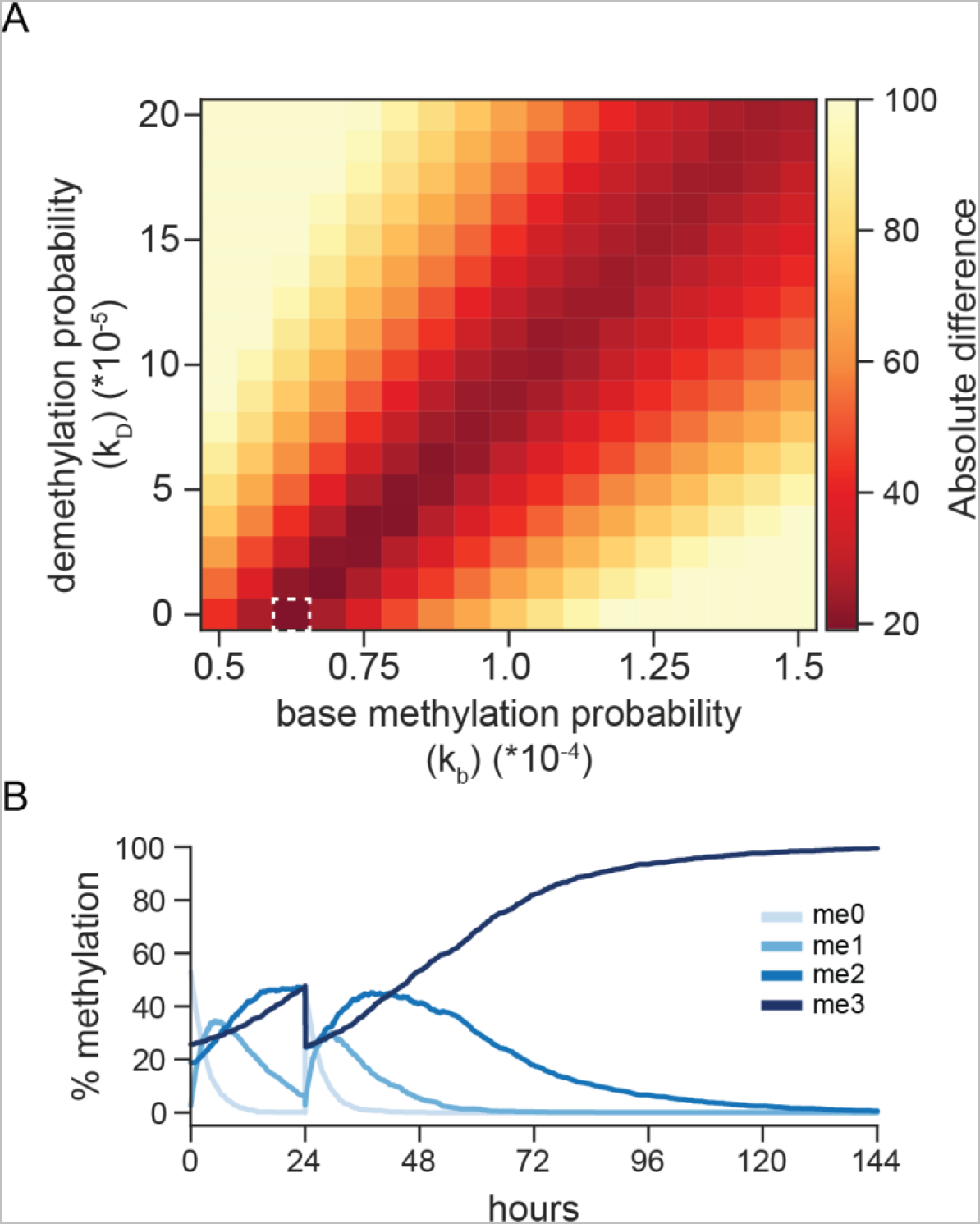
The impact of demethylation. **A**. Optimization of base methylation probability (k_b_) and demethylation probability (k_D_) parameters. The heat-map shows the sums of absolute differences between the mean fractions of un-, mono-, di-, and tri-methylated H3K27 forecasted by the model and those measured by mass-spectrometry in S2 and Kc cells for different combinations of k_b_ and k_D_ and S_3_ = 3.5. It indicates that parameter setting k_b_ = 0.625×10^-4^ and k_D_ = 0 (marked with dashed white box) result in predictions most closely resembling the experimental measurements. The fractions of variously methylated H3K27 positions predicted by the model were calculated based on 10 inspection rounds from 1000 parallel simulations. **B.** The dynamics of H3K27 methylation states over time in a representative simulation with optimal parameters (S_3_ = 3.5; k_b_ = 0.625×10^-4^; k_D_ = 0) after the replication ceased to happen.

The situation changes dramatically for terminally differentiated cells that stop replicating their DNA. Removing chromatin replication from the simulations leads to a gradual build-up of tri-methylated H3K27. As illustrated by Figure 5B, in the absence of demethylation, 95% of all histone H3 in *Drosophila* cultured cells will be tri-methylated within approximately 78 hours after the DNA replication stops. In these non-proliferating cells, this estimate assumes that the effective concentration of PRC2 remains unchanged (i.e., S_3_ = 3.5; k_b_ = 0.625×10^-4^). However, even if the effective PRC2 concentration is reduced, all H3 molecules will be converted to H3K27me3 as long as some H3K27 methylation activity is present within the cell but methyl groups are not removed. To summarize, the role of replication-independent H3K27 demethylation in proliferating cells is likely limited to specific and/or transcriptionally active loci. Yet, it is indispensable to prevent the indiscriminate H3K27 tri-methylation in cells that exited the cell cycle.

### Intergenerational inheritance of H3K27me3

As discussed above, the chromatin replication frequency has a large impact on H3K27 methylation levels. Remarkably, *Drosophila* development starts with thirteen very rapid mitotic divisions, of which the first nine each take 10 minutes or less. The first extended cell cycle (cycle 14) occurs two hours after fertilization when zygotic genes start to be transcribed. Initially, the paternal genome mostly lacks methylated H3K27, as protamines replace most histones during sperm differentiation (*53*). In contrast, the maternal genome retains nucleosomes and displays abundant H3K27me3 in the oocyte and in the maternal pronucleus just after fertilization (*54*). During the oocyte maturation the mother deposits PRC2 components in the embryo. Disrupting this supply is lethal and leads to erroneous expression of homeotic genes (*54*), the most appreciated targets of the Polycomb system. Resupplying PRC2 after the zygotic genome activation does not alleviate the problem (*54*). From this, it was proposed that intergenerationally inherited H3K27me3 is essential for the repression of homeotic genes later in development.

This conclusion is surprising in two ways. First, it implies that maternally inherited H3K27me3 survives the “dilution” during the early rapid replication cycles. Second, it suggests that the maternal genome is imprinted for the early onset and/or more efficient repression, compared to the paternal one. Intrigued by these questions, we sought to address them using our model. To this end, we simulated H3K27 methylation throughout the first 4 hours of the *Drosophila* development. We used the cell cycle timing from Ashburner and co-authors (*55*) and, for simplicity, assumed that the replication events occurred in the middle of each cycle. In a pilot attempt, we performed simulations of hit-and-run methylation with parameters optimized for cultured *Drosophila* cells (i.e., S_3_ = 3.5; k_b_ = 0.625×10^-4^) and compared the overall levels of variously methylated H3K27 forecasted by the model to those determined in 2-4 hour old embryos by mass-spectrometry (*56*). It was immediately clear that with this base methylation probability, the forecasted levels of methylated H3K27 were much lower than the ones determined experimentally (Figure 6A), because the methylation could not keep pace with rapid influx of unmodified histones (Figure 6B).

**Figure 6.**
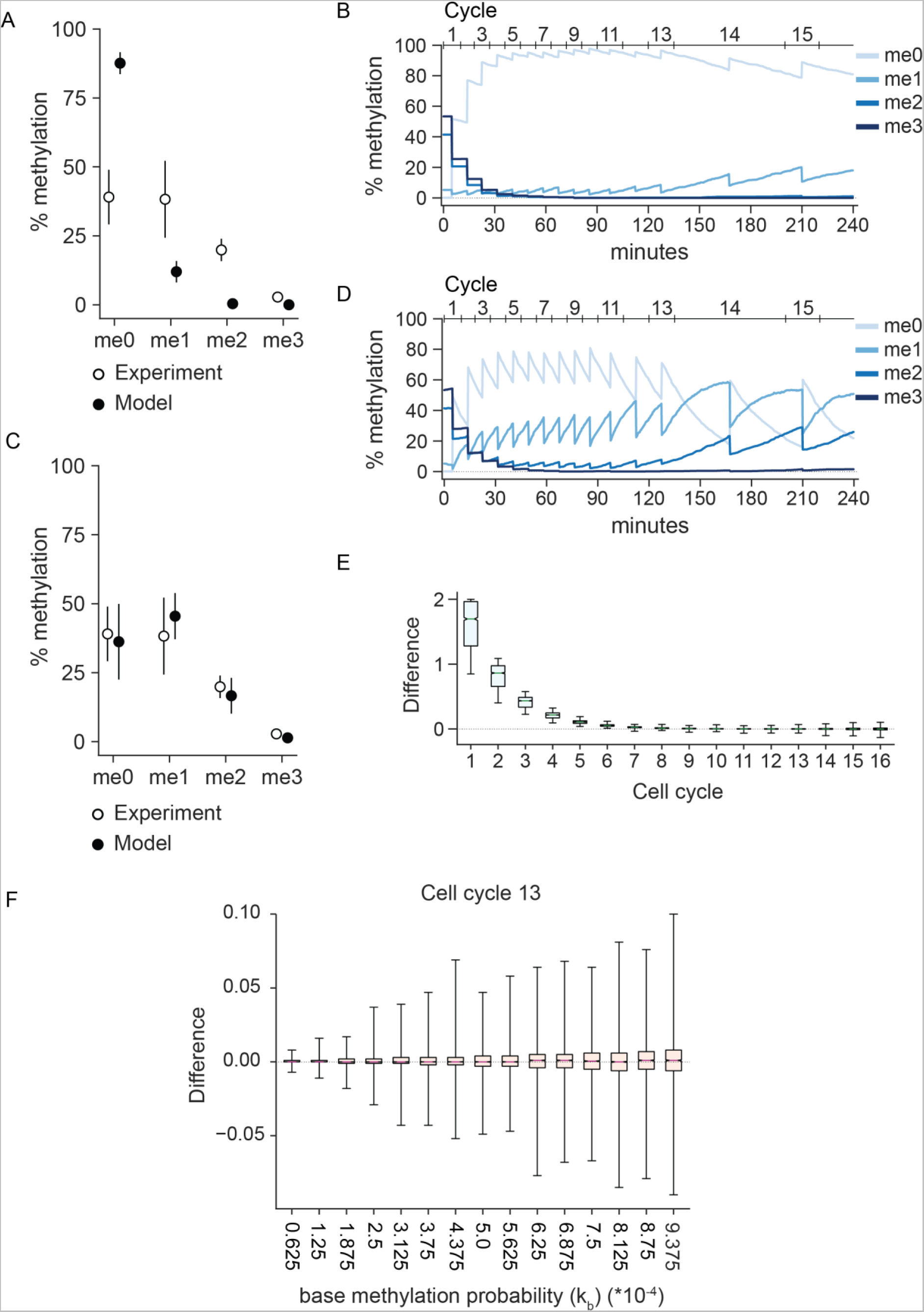
Dynamics of H3K27 methylation in developing *Drosophila* embryo. **A**. Quantifications of H3K27 methylation levels in the 2-to-4-hour embryo. The black dots indicate mean values from one random inspection taken from 1000 simulations with S_3_ = 3.5; k_b_ = 0.625×10^-4^; k_D_ = 0 parameter settings. The white dots are mean values from experimental measurements by (*56*). The bars indicate standard deviations**. B.** The dynamics of H3K27 methylation levels from a representative simulation of the first four hours of *Drosophila* embryonic development with parameter settings as in **A**. Quantification (**C**) and dynamics (**D**) of H3K27 methylation levels measured as in **A** and **B**, but with ten times higher base methylation probability (k_b_ = 6.25×10^-4^). **E.** Comparison of forecasted H3K27me3 within maternal (initially methylated) and paternal (initially unmethylated) alleles of the bithorax complex. Here and in **F** the box plots show distributions of differences (maternal-paternal) in mean H3K27me3 occurrence within individual nucleosomes of the bithorax complex. The mean H3K27me3 occurrence within a nucleosome was calculated from 1000 simulations. **F.** Forecasted H3K27me3 within maternal and paternal alleles of the bithorax complex becomes identical at cycle 13 regardless of base methylation probability.

While the first 13 mitotic divisions within the Drosophila embryo happen synchronously, the global synchrony ceases in the 14th cycle. The embryo divides into 25 mitotic domains with a locally synchronous mitosis (*57*). 4 hours after fertilization, cells in some of these domains enter the 16^th^ mitotic cycle while others remain in the 14^th^ cycle. To test whether the low H3K27 methylation forecasted by the model may be due to the unaccounted contribution of cells with a long 14^th^ cycle, we re-ran the simulations, setting the 14^th^ cycle to last until the end of the 4th hour. The forecasted methylation values did not change much and remained too low (Figure S2A-B). From this, we conclude that, compared to cultured cells, embryos must have a much higher effective concentration of PRC2 provided maternally (and therefore much higher base methylation probability k_b_).

To explore this conjecture, we performed a new set of simulations with ten-fold greater base methylation probability (k_b_= 6.25×10^-4^). These simulations forecasted bulk methylation levels that agree with those measured experimentally (Figure 6C-D, S2C-D). That is regardless of whether we used the cell cycle timing from Ashburner and co-authors (*55*) or with the long 14^th^ cycle. Simulations with the former timings (*55*) forecast methylation values that fit experimental observation better, so we used them for the following simulations.

To investigate the impact of maternally inherited H3K27me3, we focused on the *bithorax complex* homeotic gene cluster (Figure 3B), to parallel genetic experiments by Zenk and co-authors (*54*), and compared methylation dynamics across the cluster under two contrasting scenarios. In the first scenario, we started the simulations with the H3K27me3 level of 80%, which would exist in dividing cells just before the replication. In the second scenario, we performed the simulations with the same parameters but starting with unmethylated chromatin (a representation of the paternal chromatin). In the first scenario (maternal chromatin), the initial H3K27me3 profile rapidly declines, reaching the lowest level at cycle 10, after which it starts to build up, first very slowly and then more rapidly as cell cycles become longer (Figure 7). In the second case (paternal chromatin), the tiny H3K27me3 peaks centred on PREs appear as early as cycle 2. The methylation continues to accumulate with a pace that correlates with the length of the cell cycle (Figure 7).

**Figure 7.**
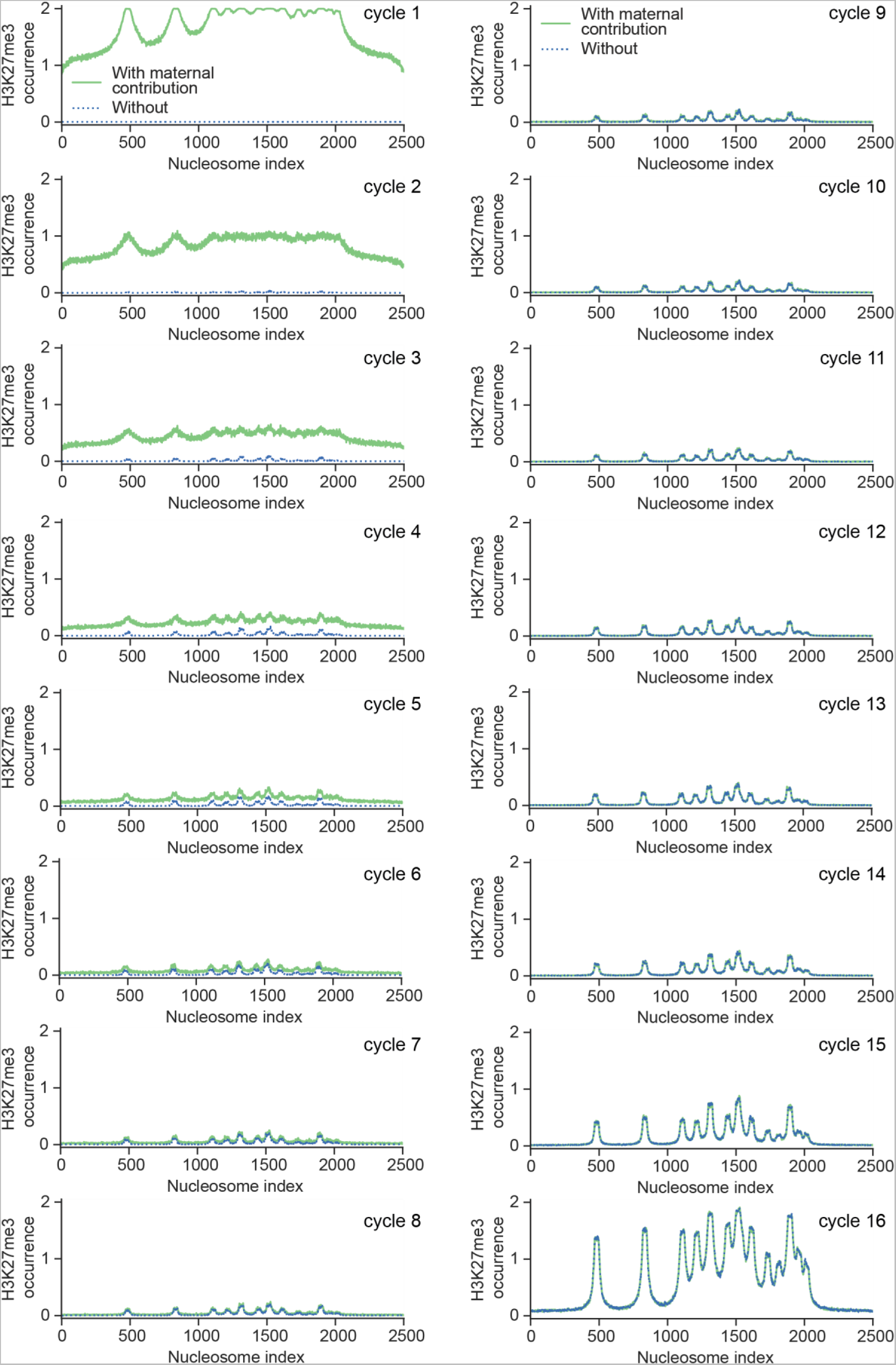
H3K27me3 profiles over the maternal and paternal alleles of the bithorax complex. The two profiles converge rapidly and become identical by cell cycle 13. The plots show mean H3K27me3 occurrence calculated from 1000 simulations with S_3_ = 3.5; k_b_ = 6.25×10^-4^.

Importantly, the H3K27me3 profiles forecasted for the two scenarios converge rapidly (as a single exponential) (Figure 6E). By cycle 13, they become identical, regardless of how much PRC2 is contributed maternally (Figure 6F). At cycle 14 – when the zygotic genome starts to be used – the maternally deposited H3K27me3 is therefore too dilute to have any impact on the repression by the Polycomb system. Even at mitotic cycle 10, when the first few zygotic genes start to be transcribed (*58*), the median difference between the number of maternal and paternal H3 molecules tri-methylated at K27 over the *bithorax complex* is only 0.003 (Figure S3). In other words, only 3 out of 1000 H3 molecules within the *bithorax complex* typically carry K27me3 inherited from the mother.

To summarize, our observations suggest that the large quantity of PRC2 deposited in the oocyte by the mother serves to attain a high effective concentration of the enzyme in the beginning of embryonic development. This is necessary to counteract rapid chromatin replication cycles and methylate the PRE-equipped loci in time, rather than to propagate the maternally inherited H3K27me3.

## Discussion

Four main conclusions can be drawn from the work presented here. First, our understanding of the Polycomb system has reached the level required to forecast PRC2-mediated histone methylation by numerical modelling. Our forecasts have high temporal resolution and spatial accuracy that matches contemporary experimental techniques. Second, the allosteric stimulation of PRC2 by the pre-existing H3K27me3 is necessary for broad and timely methylation of H3K27 around PREs. However, it is not sufficient to sustain high-level H3K27me3 if PREs are removed. Third, the methylating activity of PRC2 and the influx of unmodified histone molecules during chromatin replication balance each other to define the levels of methylated H3K27 in dividing cells. In these cells, the role of replication-independent H3K27 demethylation is likely limited to specific loci. However, the demethylation is indispensable in fully differentiated cells to prevent the indiscriminate H3K27 tri-methylation. Fourth, the intergenerationally inherited H3K27me3 does not “survive” rapid cycles of chromatin replication during *Drosophila* embryonic development and is therefore unlikely to transmit the memory of epigenetic repression to the offspring. Below we discuss these conclusions and their implications in greater detail.

H3K27 methylation by *Drosophila* PRC2 can be forecasted by a relatively simple computational framework. Three insights were important for its successful implementation. We realized that the bulk levels of variously methylated H3K27, measured experimentally by others (*41, 56*), are produced by hit-and-run action that resembles the PRC2-mediated methylation reaction *in vitro*. We recognized that accounting for chromatin replication provides a simple way to introduce time in a purely probabilistic Monte-Carlo model. Finally, we correctly guessed that a calibrated hit-and-run model could be adapted to represent H3K27 methylation at PRE-equipped genes by converting the degree of PRC2 tethering by a PRE into a higher methylation probability.

Animal cells produce several variants of PRC2 (*59*). All of them contain the same catalytic core but differ in several mutually exclusive subunits. The contribution of distinct PRC2 complexes to H3K27 methylation at PRE-equipped genes relative to that genome-wide is not fully understood. It is likely that certain variant complexes are tethered to PREs better than others and that the variant PRC2 complexes differ in their catalytic efficiency. Nevertheless, our modelling suggests that just tethering PRC2 from the nucleoplasm to a specific locus is, in principle, sufficient to produce H3K27me3-enriched chromatin domains over the loci repressed by the Polycomb system.

The allosteric stimulation by the pre-existing H3K27me3 is a peculiar property of PRC2 and the Polycomb system does not function in *Drosophila* mutants with impaired stimulation (*23*). It would be tempting to imagine that tethering PRC2 is necessary only to establish the high degree of H3K27 tri-methylation around the PRE, and that allosteric stimulation of untethered PRC2 is sufficient to renew it after the DNA replication and cell division. Still, experiments with *Drosophila* transgenes equipped with one PRE showed that H3K27me3 is rapidly lost from the transgene when the PRE is excised by recombination (*9, 10*). Our simulations indicate that H3K27 methylation decays when removing PREs from any locus, regardless of the number of PREs or their relative arrangement. Furthermore, they show that the allosteric stimulation likely is relatively unimportant for the bulk H3K27 methylation by untethered PRC2, because the average genomic density of H3K27 tri-methylated nucleosomes remains too low for most of the cell cycle. Instead, the simulations indicate that the stimulation of PRC2 by H3K27me3 enables broad H3K27 tri-methylation around PREs and increases the speed with which the H3K27me3 is regained after chromatin replication. To summarize, the allosteric stimulation makes PRC2 better in methylating H3K27 at sites where it is needed the most (i.e. PRE-equipped developmental genes) as opposed to indiscriminate methylation of the entire transcriptionally inactive genome.

Our simulations show that, in dividing cells, the pools of methylated H3K27, overall and at PRE-equipped genes, are defined by the balance between the methylating activity of PRC2 and the influx of unmodified H3 molecules during chromatin replication. The two major opposing factors that set the balance are the effective concentration of PRC2 in the nucleus and the replication frequency. In line with this notion, experiments with cultured cells homozygous for a temperature-sensitive mutation in the *E(z)* gene indicate that the replication-independent loss of H3K27 methylation across the genome is slow (*17*).

Mutations affecting PRC2 activity, such as gain-of-function, hypomorph and loss-of-function mutations in genes encoding the core components, are widespread in cancer (*60*). Several PRC2 inhibitors were shown to impair the proliferation of cultured cancer cells and are now in clinical trials (*61*). Lowering PRC2 activity is expected to alter the balance between methylation of H3K27 and the influx of unmodified histone molecules. This balance shifts even more in highly proliferating cells. As follows from our simulations, a treatment with a small molecule that stimulates PRC2 activity will also tilt the balance but in the opposite direction. In cancer cells with hyperactive gain-of-function PRC2 mutations, this may lead to indiscriminate tri-methylation of H3K27 throughout the genome and cell death. This counterintuitive strategy has not yet been tried.

There are several examples when a regulatory state of a gene is passed from parents to their offspring (*62, 63*), including that of transgenic *Drosophila* epialleles repressed by the Polycomb system (*64*). PRC2 components are deposited in *Drosophila* embryos by the mother during oogenesis. The maternal *Drosophila* genome retains nucleosomes and displays abundant H3K27me3 in the oocyte or in the maternal pronucleus just after fertilization and the loss of the maternal PRC2 supply is lethal (*54*). From this, it was proposed that intergenerationally inherited H3K27me3 is critical for *Drosophila* embryonic development. Our simulations do not support this hypothesis. They argue that intergenerationally inherited H3K27me3 does not survive the rapid replication cycles during *Drosophila* embryonic development and therefore are unlikely to directly transmit the memory of epigenetic repression to the offspring. Instead, it appears that high amounts of maternally provided PRC2 are needed to enable timely methylation of PRE-equipped loci when the zygotic genome activation occurs. Our observations do not exclude a role for the intergenerationally inherited H3K27me3 in organisms with slower development. The first cell divisions within mouse or human embryos take approximately 12 hours. Remarkably, recent reports suggest that in very early mouse and human embryos a small number of maternally imprinted genes lack DNA methylation but harbour maternal allele-specific H3K27me3 required to maintain the inactive state of these genes until the implantation stage (*65–67*).

To conclude, we developed a framework to forecast the PRC2-catalyzed methylation in *Drosophila* because, in this organism, the PRE locations, the extent of PRC2 tethering by the individual elements, and the bulk levels of variously methylated H3K27 have been reliably measured. With rapid progress in understanding PRC2 tethering to other genomes, we foresee that our model will be adaptable to forecast H3K27 methylation in broad range of organisms, including mice and humans.

## Acknowledgements

We are grateful to Dr. Jan Larsson and Dr. Tatyana Kahn for critical comments to the manuscript. This work was supported by grants from Swedish Research Council to Y.B.S. (2021-04435) and L.L. (2021-04080) and Cancerfonden to Y.B.S (22 2285 Pj). This research was conducted using the resources of High Performance Computing Center North (HPC2N), provided by the Swedish National Infrastructure for Computing (SNIC), partially funded by the Swedish Research Council through grant agreement no. 2018-05973.

## Supplementary figures

**Figure S1.**
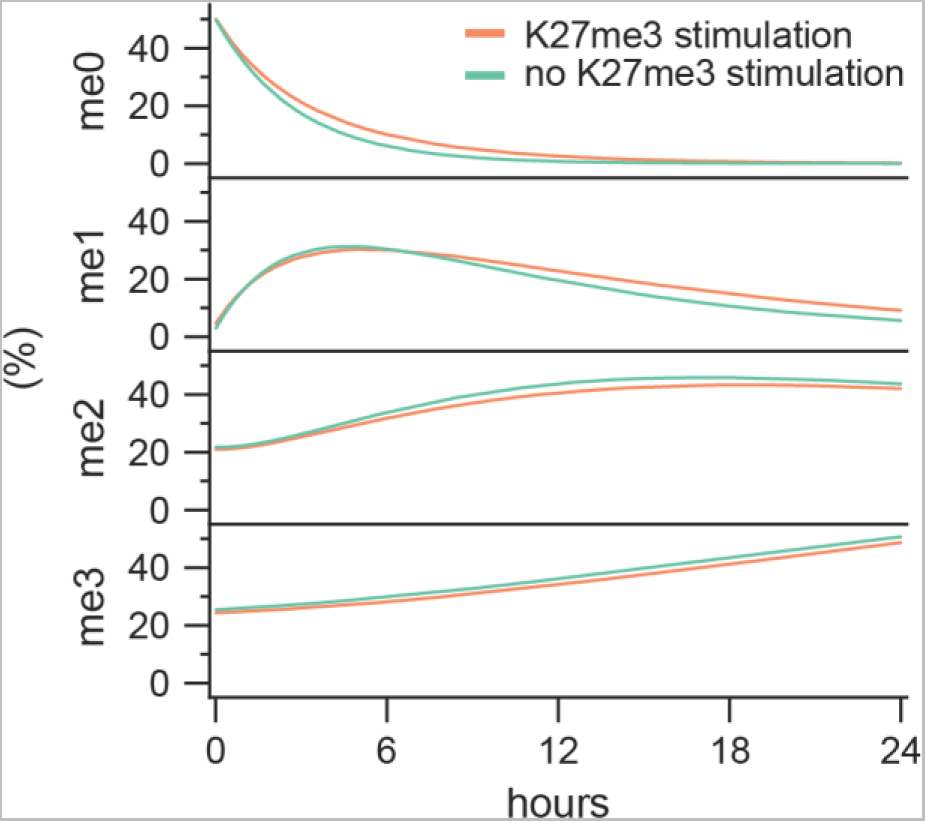
The impact of PRC2 stimulation by H3K27me3 on hit-and-run methylation dynamics. Forecasted relative abundance of variously methylated H3K27 over one cell cycle in the presence (orange, S_3_ = 3.5 and k_b_ = 0.625×10^-4^) or in the absence (green, S_3_ = 1 and k_b_ = 1.125×10^-4^) of H3K27me3 stimulation. The graphs show mean values over 1000 parallel simulations with shaded area corresponding to 95% CI.

**Figure S2.**
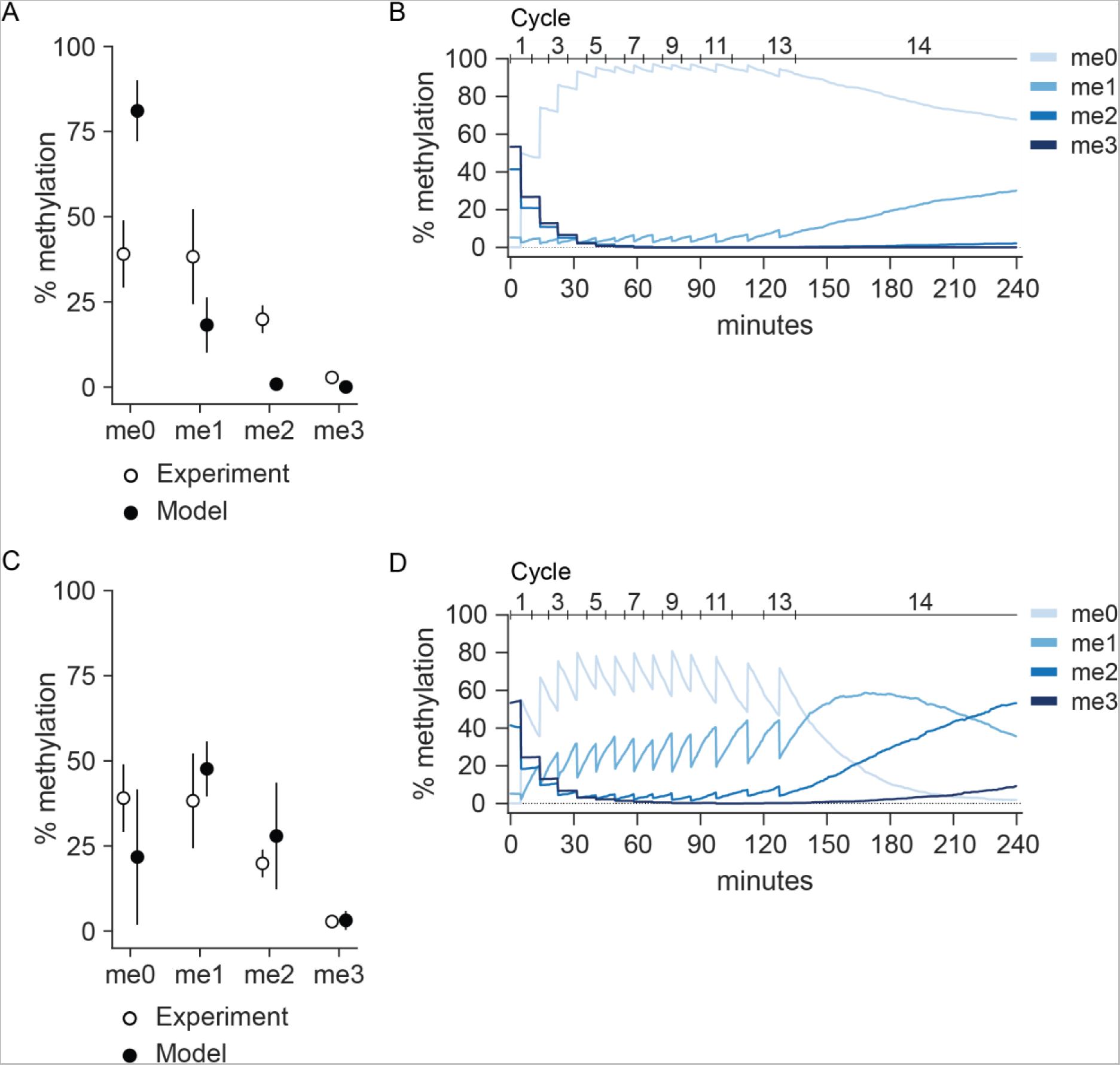
Dynamics of H3K27 methylation in developing *Drosophila* embryo assuming long 14^th^ mitotic cycle. **A**. Quantifications of H3K27 methylation levels in the 2-to-4-hour embryo. The black dots indicate mean values from one random inspection taken from 1000 simulations with S_3_ = 3.5; k_b_ = 0.625×10^-4^; k_D_ = 0 parameter settings. The white dots are mean values from experimental measurements by (*56*). The bars indicate standard deviations**. B.** The dynamics of H3K27 methylation levels from a representative simulation of the first four hours of *Drosophila* embryonic development with parameter settings as in **A**. Quantifications (**C**) and dynamics (**D**) of H3K27 methylation levels measured as in **A** and **B**, but with ten times higher base methylation probability (k_b_ = 6.25×10^-4^).

**Figure S3.**
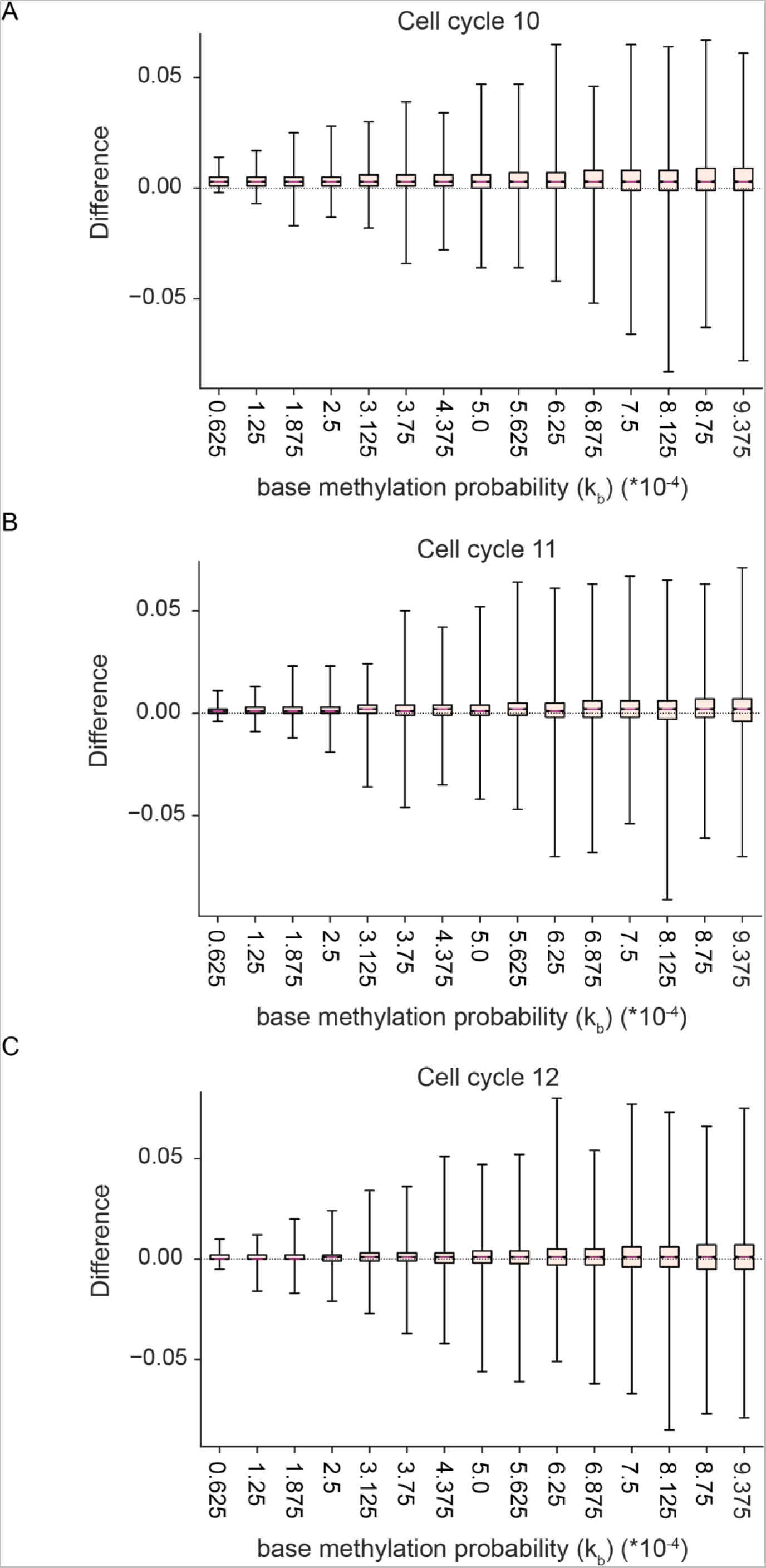
Forecasted H3K27me3 within maternal and paternal alleles of the bithorax complex converge equally fast at various base methylation probabilities. Comparison of forecasted H3K27me3 within maternal (initially methylated) and paternal (initially unmethylated) alleles of the bithorax complex during cell cycles 10 (**A**), 11 (**B**) and 12 (**C**). The box plots show distributions of differences (maternal-paternal) in mean H3K27me3 occurrence within individual nucleosomes of the bithorax complex. The mean H3K27me3 occurrence within a nucleosome was calculated from 1000 simulations.

